# Hypergraphs for predicting essential genes using multiprotein complex data

**DOI:** 10.1101/2020.04.03.023937

**Authors:** Florian Klimm, Charlotte M. Deane, Gesine Reinert

**Affiliations:** Department of Statistics, University of Oxford, Oxford OX1 3LB, United Kingdom; Department of Mathematics, Imperial College London, London, SW7 2AZ, United Kingdom; MRC Mitochondrial Biology Unit, University of Cambridge, Cambridge Biomedical Campus Hills Road, Cambridge, CB2 0XY, United Kingdom

**Keywords:** Gene essentiality, Protein interaction networks, Hypergraphs, Null models, Hierarchical exponent, Centrality, Clustering coefficient

## Abstract

Protein-protein interactions are crucial in many biological pathways and facilitate cellular function. Investigating these interactions as a graph of pairwise interactions can help to gain a systemic understanding of cellular processes. It is known, however, that proteins interact with each other not exclusively in pairs but also in polyadic interactions and they can form *multiprotein complexes*, which are stable interactions between multiple proteins. In this manuscript, we use *hypergraphs* to investigate multiprotein complex data. We investigate two random null models to test which hypergraph properties occur as a consequence of constraints, such as the size and the number of multiprotein complexes. We find that assortativity, the number of connected components, and clustering differ from the data to these null models. Our main finding is that projecting a hypergraph of polyadic interactions onto a graph of pairwise interactions leads to the identification of different proteins as hubs than the hyper-graph. We find in our data set that the hypergraph degree is a more accurate predictor for gene-essentiality than the degree in the pairwise graph. We find that analysing a hypergraph as pairwise graph drastically changes the distribution of the local clustering coefficient. Furthermore, using a pairwise interaction representing multiprotein complex data may lead to a spurious hierarchical structure, which is not observed in the hypergraph. Hence, we illustrate that hypergraphs can be more suitable than pairwise graphs for the analysis of multiprotein complex data.

## 1 Introduction

Protein-protein interactions represent the chemical reactions and physical contacts between proteins [1]. Their statistical analysis can give insights into underlying cellular processes and the organism they govern. They are therefore used in various bioinformatics applications, such as, the reconstruction of phylogenetic trees, the prediction of proteins’ biological functions and the identification of functional modules (for reviews see [1, 2]). One important application is the prediction of whether a gene that codes a certain protein is essential [3, 4]. Typically, a dataset of protein-protein interactions is represented as a binary undirected network, with proteins as nodes and edges representing interactions. For predicting essential proteins, one can for example, investigate the centralities of nodes [5] or combinations of multiple measures [6] in such a protein-protein interaction network.

In such studies, the interaction between the proteins are modelled as pair-wise. More than half of all proteins, however, form *multiprotein complexes* that may consist of more than two proteins that are linked by non-covalent interactions [7, 8]. Protein complexes are crucial for most biological processes *ATP synthase*, for example, an enzyme that creates the energy storage molecule adenosine triphosphate (ATP), consists of up to eight different subunits, each a protein [9]. Proteins can be involved in different complexes or have additional activities, independent of the complex itself [10]. These biological observations indicate that mathematical objects that take multiprotein complex information as high-order interactions into account might be an appropriate way to study cellular systems in general and the prediction of essentiality, specifically. In this study, we use *hypergraphs* [11], which are one way to represent *polyadic interactions* (i.e., interactions of higher order than pairwise), to analyse a network of human multiprotein complexes.

In Fig. 1, we give three examples of multiprotein complexes and the representation of their high-order interactions as *hyperedges* in a hypergraph. The *exon junction complex* is a crucial molecular machine that influences the translation of mRNA molecules [12]. It consists of four different protein components CASC, Y14, MAGOH, and EIF4A3 and we therefore represent this interaction as a four-edge in a hypergraph. Two of these four proteins (Y14 and MAGOH) can form a separate complex (the Y14–MAGOH complex), which we represent as a two-edge [13]. The PYM protein can bind to this complex and we represent the formed complex which consists of three proteins as a three-edge. These three different complexes demonstrate only a small subset of the complex higher-order interactions that we observe in the human body. The PYM protein, for example, interacts with the 40S ribosomal subunit, which itself consists of thirty-three proteins (not shown). Higher-order interactions are common in cellular processes and in this study, we represent their complex interaction structure as a hypergraph.

**Figure 1:**
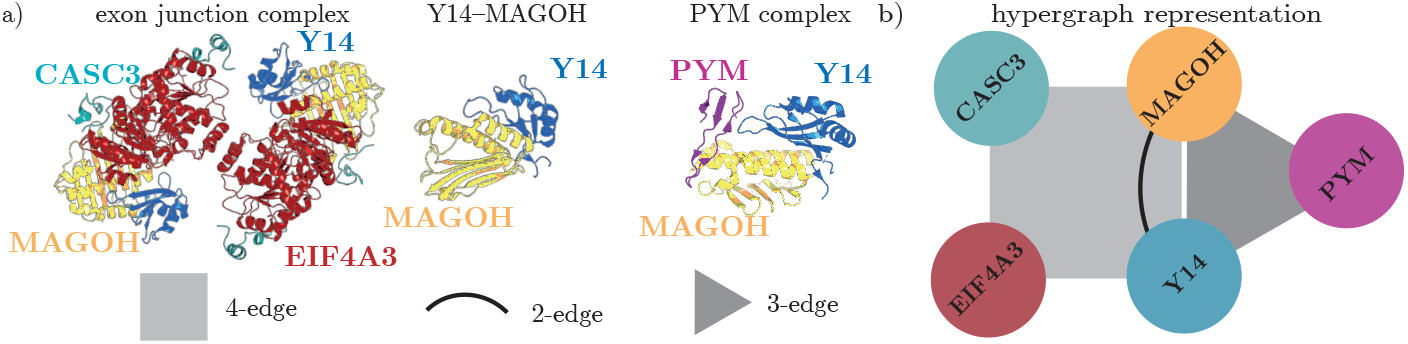
Protein may interact with each other and form complexes. (a) We show the exon junction complex, the Y14–MAGOH complex, and the PYM complex in cartoon representations. We can represent these interactions as hyperedges, whose cardinality is the number of different involved proteins. The exon junction complex, for example, consists of four different proteins (CASC3, Y14, MAGOH, and EIF4A3; shown in green, blue, yellow, and red, respectively) and thus we represent it as a four-edge. The two proteins Y14 and MAGOH can also interact with each other and we represent this interaction as a 2-edge. The interaction between PYM, Y14, and MAGOH is represented in a 3-edge. (b) We jointly represent these three interactions in a hypergraph, with the *N* = 5 nodes representing proteins and the *M* = 3 hyperedges representing multiprotein complexes.

Other mathematical structures are potentially also suited to represent high-order interactions. *Simplicial complexes*, for example, have been used to investigate time-series [14, 15], and many other systems [16, 17, 18]. For our purposes, we argue that hypergraphs are a more suited mathematical framework because simplicial complexes require *set inclusion*^1^, which for our application implied that for every multiprotein complex, all subsets of constituent proteins would also form a multiprotein complex, which in general is not the case.

Hypergraphs have been identified as a framework to investigate metabolic pathways, which describe molecular reactions [20, 21, 22] and some methods for the statistical analysis of hypergraphs have been developed (e.g., centralities [23], local clustering coefficient [21], configuration models [24]). In this manuscript, we focus on the degree and clustering coefficient as node-measures because they are commonly used for predicting the lethality or essentiality of proteins [3, 5].

First, we assess whether the constructed hypergraph contains signal beyond the degree. To test this, we construct two random null models, one an existing hypergraph configuration model [24] and one a new Erdős-Rényi-type hyper-graph model. We compare the hypergraph properties of these null models with the data hypergraph. Similarly to many empirical graphs, we find that assortativity, clustering, and number of connected components differ strongly from these random null models.

Often, one projects higher-order interactions to pairwise interactions to be able to use a broad selection of tools that have been developed for the analysis of graphs. To test which representation, graph or hypergraph, is more suitable for the analysis of the multiprotein complex data, we compare gene essentiality data from the *Online GEne Essentiality database* [25] with the degree in the hypergraph and the degree in the pairwise interaction graph. As the former is in stronger agreement, the hypergraph representation outperforms the pairwise graph representation for identifying essential genes using degrees.

Next, we show that using a pairwise interaction graph may lead to a spurious result for the commonly asserted *hierarchical organisation* of complex systems: together with the degree, the local clustering coefficient has been used to quantify the hierarchical organisation of pairwise complex networks [26, 27] in general and metabolic networks, specifically [28, 29]. Here, we demonstrate that projecting a hypergraph onto a graph can drastically increases the local clustering coefficient of many nodes. Furthermore, this projection may indicate a statistically significant hierarchical organisation of the graph that is not observed in its hypergraph form. As such a projection is common in many network studies —either explicitly or implicitly— one should be careful about the interpretation of such results.

Overall, we propose that a hypergraph representation for multiprotein complex data is a better approach to identify essential genes and that not using this representation may lead to a spurious hierarchical structure in the graph.

## 2 Methods

### 2.1 Hypergraph measures

A *graph* is an ordered pair 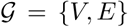, where *V* is a set of nodes and *E* ⊂ *V × V* a set of edges that connect the nodes pairwise; in this paper, edges are undirected and unweighted. Two nodes which are connected by an edge are called *neighbours* and the *neighbourhood* 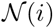 of a node *i* is the set of all its neighbours. A *hypergraph* is a generalisation of a graph that allows edges that connect more than a pair of nodes and are therefore called *hyperedges*. Formally, we define 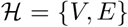, where *V* is a set of nodes and the hyperedge set *E* is a subset of the *power set P* (*V*), which is the set of all subsets of *S*, but excluding the empty set ∅. Therefore, a hyperedge may connect any set of nodes but not the empty set. The number *c* = |*e*| of nodes that a hyperedge *e* ∈ *E* connects to is called the hyperedge’s *cardinality*. A hyperedge with cardinality *c* is also called a *c-edge*.

For graphs, the degree *k*_*i*_ of a node *i* is the number of edges it connects to. For simple graphs (i.e., graphs without parallel edges and without self-loops), the degree is identical to the number of neighbours this node has. In accordance with graphs, the degree 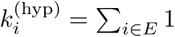 of node *i* in a hypergraph is the number of hyperedges it connects to. In contrast to simple graphs, the degree of a node in a hypergraph is not necessarily equal to the number of its neighbours [30]. The maximum degree max(*k*) is the largest degree of any node in a hypergraph.

The *local clustering coefficient C*_*i*_ of a node *i* in a graph is

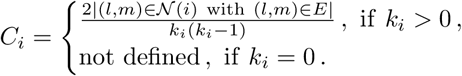

We choose a definition from [21] to generalise the *local clustering coefficient* but adapt it slightly for clarity. The local clustering coefficient 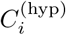 of a node *i* in a hypergraph is

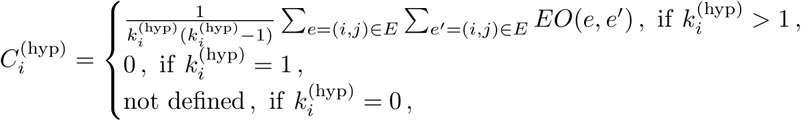

where the *extra overlap EO*(*e, e*′) between two intersecting hyperedges *e* and *e*′ is defined as

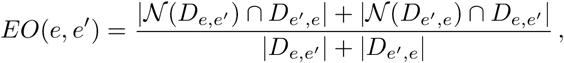

with *D*_*e,e*′_ = *e* − *e*′, the (asymmetric) set difference between *e* and *e*′. We define *EO*(*e, e*′) = 0 for *e* = *e*′. The neighbourhood 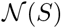 of a set *S* of nodes is the union of the neighbourhoods of each node in the set, i.e. 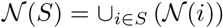. For isolated nodes with 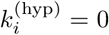, the local clustering is not defined. Another variant of local clustering is suggested in [20] and a global clustering coefficient is discussed in [23]. The mean local clustering coefficient ⟨*C*_*i*_⟩ is defined as 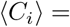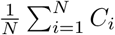. The *assortativity ρ* of a hypergraph is a measure of the correlation between a node’s degree and the degree of its neighbours. Following [24], the assortativity *ρ* of a hypergraph is the Pearson correlation between the nodes’ degrees 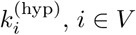 and the mean degree 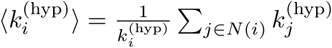 of all of its neighbours.

For graphs, the relationship between degree *k*_*i*_ and local clustering coefficient *C*_*i*_ has been approximately described through a power-law 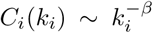 in which *β* is called the *hierarchical exponent* [26]. In practice, we estimate *β* by calculating the Pearson correlation between log_10_(*k*_*i*_) and log_10_(*C*_*i*_) for *i* = 1, …. *N*. With the definitions of local clustering coefficient and degree, we can also compute the *hypergraph hierarchical exponent β*^(hyp)^ as Pearson correlation between 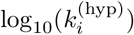 and 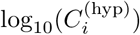 for *i* = 1, …. *N*.

For a graph, a *component* is a subgraph in which any two nodes are connected to each other by paths. For hypergraphs, more nuanced definitions exists, e.g., ‘*j*-component’ are sets of vertices such that consecutive edges in paths intersect in at least *j* vertices [31]. We discuss here exclusively 1-components and call them ‘components’ for simplicity. The number *n*_com_ of components is the amount of components in a hypergraph. The size *S*^(*m*)^ of a component is the number of nodes in it. We also compute the relative size *S*_max_/*N* ∈ (0, 1] of the largest connected component.

### 2.2 Data and preprocessing

We constructed a hypergraph from Reactome version May 2019 [32]. The data set ‘Human complexes with their participating protein molecules’ consists of a total of ~ 12, 000 complexes. The complexes include not only proteins but also other ligands, for example, small molecules (described by ‘chebi’ codes) and RNA molecules (described by ‘ensemble’ IDs). We ignored these entities and only kept entities that describe proteins with a uniprot ID. After deletion of duplicate entries, we obtained a hypergraph with *N* = 8243 nodes (representing proteins) and *M* = 6688 hyperedges (representing multiprotein complexes). This data combines obligate and non-obligate protein complexes, as well as, transient and stable protein complexes. We used gene-essentiality data from the *Online GEne Essentiality database* (OGEE) v2 [25]. We mapped genes to proteins with the Retrieve/ID mapping tool from UniProt [33].

### 2.3 Null models

Null models for hypergraphs have been developed in different domains. In uniform random hypergraphs all hyperedges have the same cardinality *c* [34, 35]. In this study, we use two different null models (see Fig. 2) that are non-uniform. For the construction of a *configuration model* of hypergraphs, we use definitions from [24]. We also define a novel null model, called Erdős-Rényi-type hypergraph model (ER-type hypergraph model), that does not fix the degrees of nodes in the hypergraph. For the configuration model, we use a randomisation algorithm. For the latter null model, in contrast, we construct hypergraphs directly.

**Figure 2:**
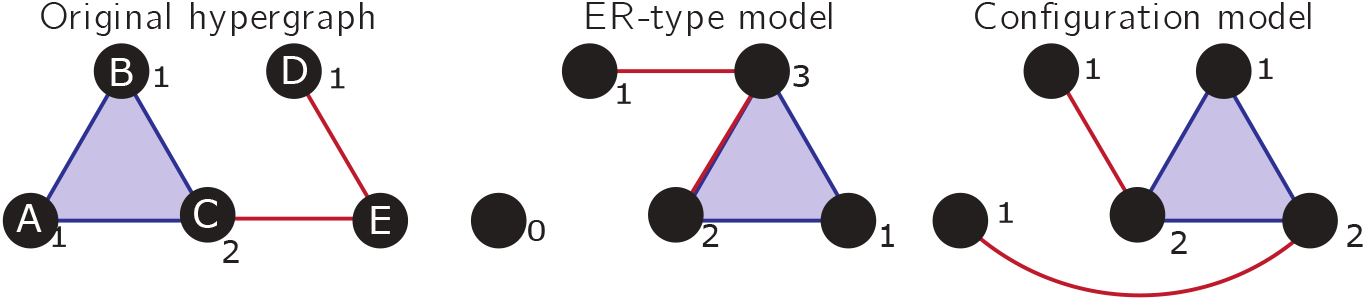
We discuss two null models. Both preserve the number *N* of nodes, the number *M* of hyperedges, and the cardinality of the hyper-edges, which in this example is (3, 2, 2). We indicate the degree of each node as number next to it. In the *degree-preserving null model* the degree of each node is preserved. In the *ER-type null model* the degree is not preserved.

#### Erdős-Rényi-type hypergraph model

For a hypergraph with *N* nodes and *M* hyperedges, we define the degree sequence **k** as the N-vector in which the *i*th element is the degree 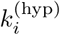 of node *i*. Similarly, we define the cardinality sequence **c** as the M-vector in which the *i*th element is the cardinality *c*_*i*_ of hyperedge *i*.

Let 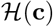 be the set of all hypergraphs with a fixed cardinality sequence **c**. The random hyperedges hypergraph model is the uniform distribution on 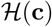 with self-loops and multiple hyperedges possible. We construct realisations of this model by connecting uniformly at random *c*_*i*_ nodes for every hyperedge *i*. In Algorithm 1, we show the procedure used to construct ER-type hypergraphs. For dense hypergraphs, this algorithm may construct hypergraphs with multiple hyperedges (i.e., hyperedges that connect the same set of nodes). For our examples, this was, however, not the case, because we do not have multiple multiprotein complexes that connect the identical proteins.

For graphs, (i.e., *c*_*i*_ = 2 for all edges) with low connection density, this random hypergraph model is identical to the *G*(*N, M*) model by Erdős-Rényi [36]. Therefore we call it *Erdős-Rényi-type hypergraph model*. This model has some similarity with the *Poisson random hypergraph* in which the number of hyper-edges between two nodes is Poisson distributed [37, 38].

#### Configuration hypergraph model

Let 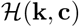 be the set of all hyper-graphs with a fixed degree sequence **k** and a fixed cardinality sequence **c**. The *vertex-labelled hypergraph configuration model* is then the uniform distribution on 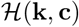. To construct random hypergraphs, we use a pairwise reshuffeling algorithm [24], which preserves the degree sequence and the cardinality sequence. As the reshuffling algorithm is an irreducible, reversible, and aperiodic Markov Chain, it has an equilibrium distribution, which we call the configuration hypergraph model. In this model, self-loops and multiple hyperedges are possible.

### 2.4 Constructing graphs from hypergraphs

We construct a *representing graph* from a hypergraphs as follows. The *representing graph R*(*H*) = (*V*′, *E*′) of a hypergraph *H* = (*V, E*) is the graph with the same set *V*′ = *E*′ of vertices as the hypergraph, and edges between all pairs of vertices contained in the same hyperedge (i.e, (*i, j*) ∈ *E*′ if there exists an edge *e* ∈ *E* such that (*i, j*) ⊂ *e*). Thus, in the representing graph, hyperedges are translated into complete subgraphs of simple edges.

**Algorithm 1:**
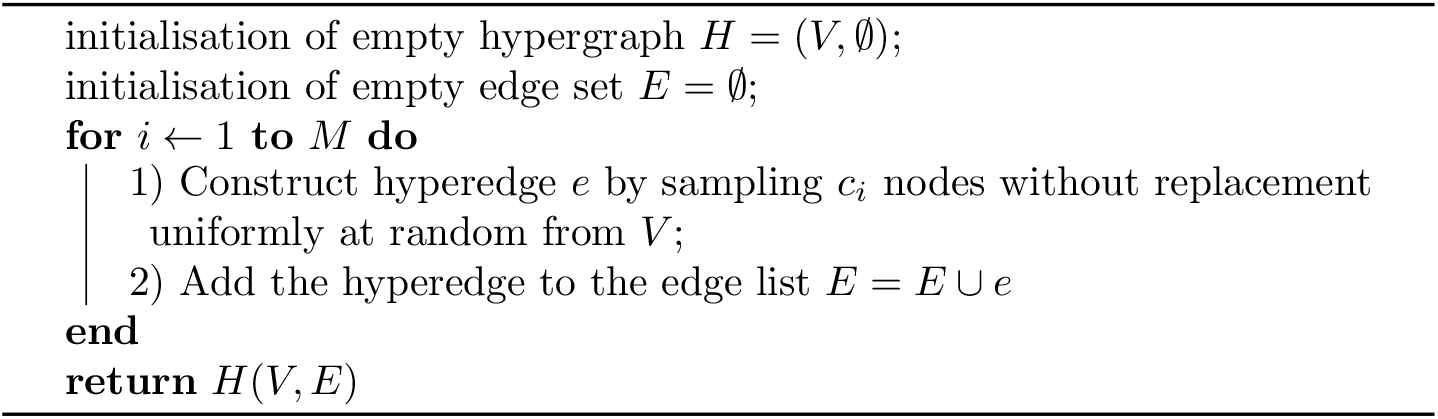
Constructing an Erdős-Rényi-type hypergraph *H* with *N* nodes and a cardinality sequence **c** = (*c*_1_, *c*_2_, … , *c*_*M*_)

**Figure 3:**
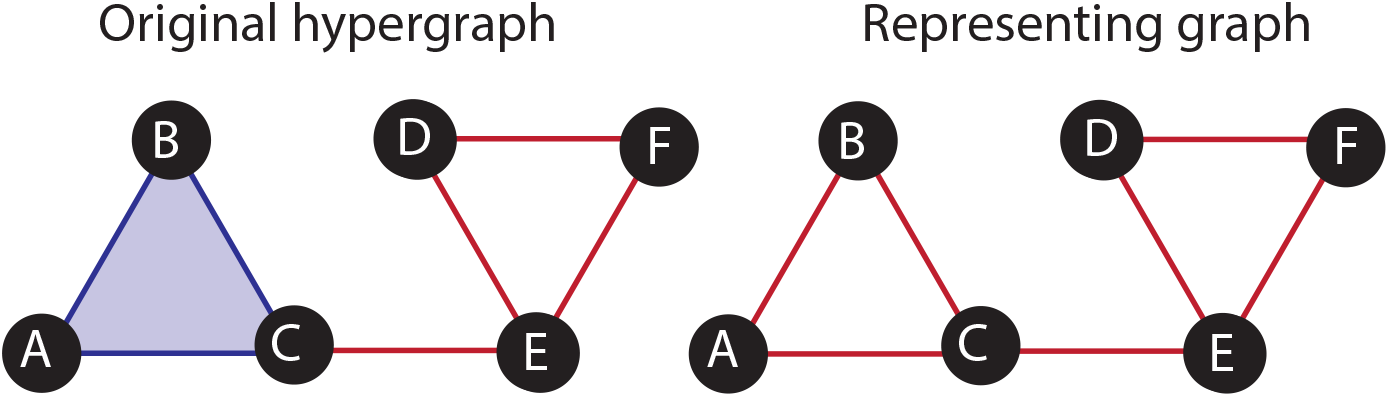
To construct the *representing graph* of a hypergaph, we replace every hyperedge of cardinality *c* with *c*(*c* − 1)/2 2-edges.

Alternatively, one could construct the *dual graph* from the hypergraphs. This method is discussed in SI A.

## 3 Results

### 3.1 Random topology null models

First, we explore whether the multiprotein complex hypergraph can be modelled (a) by the ER-type hypergraph model, which uses only its cardinality sequence **c** or (b) by the configuration hypergraph model, which uses only the hypergraph’s degree sequence **k** and cardinality sequence **c** (see Subsection 2.3 for definitions of both models). For both null models, we construct 100 independent realisations. We focus on graph measures (maximum degree, degree-assortativity, number of components, relative size of the largest component, and the mean local clustering); Table 1 gives their mean and standard deviation in these simulated hypergraphs, and the values for the data hypergraph. In SI C, we illustrate the distributions of these measures for the simulated hypergraphs.

The maximum degree in the multiprotein hypergraph is max(*k*_*i*_) = 283; in the ER-like hypergraph the maximum degree max(*k*_*i*_) ≈ 27 is far smaller. By contrast, the maximum degree max(*k*_*i*_) = 283 is by construction the same in the configuration model and the data. Computing the degree assortativity for the hypergraph yields *ρ*_data_ ≈ 0.44. This indicates that proteins with high degree *k*_*i*_ tend to form complexes with other proteins with high degree [24]. For both null models, we find a assortativity close to zero. Performing a Monte-Carlo test yields a p-values *p* < 0.01 (see Fig. 12 in SI D), indicating that assortativity of the multiprotein hypergraph is significantly larger than expected by either null model.

**Table 1:** Structural information about the protein hypergraph and the two investigated null models (ER-type model and configuration model). We constructed 100 null models and present the mean ± standard deviation. For definitions of hypergraphs measures see Subsection 2.1.

|  | original<br>hypergraph | ER-type<br>model | configuration<br>model |
| --- | --- | --- | --- |
| number of nodes | 8243 | 8243 | 8243 |
| number of hyperedges | 6688 | 6688 | 6688 |
| maximum degree | 283 | $21.3 \pm 1.4$ | 283 |
| degree-assortativity | 0.44 | $0.000 \pm 0.008$ | $-0.005 \pm 0.007$ |
| number of components | 253 | $1.01 \pm 0.01$ | $3.69 \pm 1.5$ |
| size of largest component | 0.8794 | $1 \pm 0$ | $0.9993 \pm 0.0004$ |
| mean<sup>2</sup> local clustering | 0.079 | $0.39 \pm 0.0008$ | $0.48 \pm 0.001$ |

For the mean local clustering ⟨*C*_*i*_⟩, we find that the original hypergraph has a significantly smaller clustering than both, the ER-type model and the configuration model. This contrasts with pairwise protein interaction networks, in which the clustering is normally higher than in random null models [39].

We next investigated the components of the hypergraph. The hypergraph has *n*_com_ = 253 components of which the two largest consist of 7249 and 93 nodes, respectively. This means that almost 88 % of all nodes belong to the largest component. The smaller components range in size from 22 to 2. There are 131 components which have the minimum size 2 and thus are two proteins that are connected by a 2-edges and otherwise not involved in a multiprotein complex.

In both null models, the number *n*_com_ of connected components is much smaller than in the data hypergraph. For the ER-type model and the configuration model, we obtain *n*_com_ = 1.01 ± 0.01 and *n*_com_ = 3.69 ± 1.5, respectively. This indicates that if the edges were evenly distributed between the nodes, there would exist a path between almost all nodes. Fixing the degree distribution of the hypergraph leads to a slightly larger number of connected components. This occurs because of the larger number of nodes with degree *k*_*i*_ = 1 than in the ER-like model which has a mean degree of ⟨*k*_*i*_⟩ ≈ 1.6. The relative size *S*_*max*_/*N* of the largest connected component is 1 ± 0 for the ER-type model and 0.9993 ± 0.0004 for the configuration model. For both null models, the number *n*_com_ of connected components is significantly smaller and the size *S*_max_/*N* of the largest connected component is significantly larger than for the protein data (p-values *p* < 0.01 shown in Fig. 13 in SI D).

### 3.2 Degree distribution in graph and hypergraph

Next, we compare the degrees of the nodes in the hypergraph with the degree of nodes in the representing graph. To construct the representing graph, we replace each hyperedge of cardinality *c* with *c*(*c* − 1)/2 simple edges. Therefore, the total number of edges in the representing graph is at least as large as the number of hyperedges in the hypergraph, and the degree distribution is wider for the representing graph (see Fig. 4). This indicates that replacing higher-order interactions with pairwise edges broadens the degree distribution. The mean graph-degree 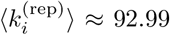 is an order of magnitude larger than that of the hypergraph 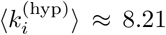. In the right panel of Fig. 4, we plot the hypergraph-degree with the degree in the representing graph for each protein. The Spearman correlation between both is 0.34, which indicates a weak correlation between the two quantities. The genes with highest degree, RPS27A and UBA52, are the same in both structures. These genes encode the protein ubiquitin, which targets proteins to degrade them and is known to bind to many different proteins [40]. For the hypergraph degree 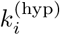, Ubiquitin B (UBB) has the third highest degree. For the representing graph degree 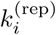, GNB1 and GNGT1, two guanine nucleotide-binding proteins have the third and fourth highest degrees. Both proteins are membrane-bound proteins that form complexes consisting of a large number of proteins. In Fig. 5, we show a force-directed layout of the representing graph. The size of the nodes is proportional to their representing graph degree 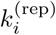 and the colour indicates the hypergraph degree 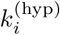. We observe cliques of nodes with high 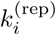 and low 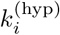: these nodes represent proteins that participate in a large multiprotein complex but no other interactions. These observations indicate that representing multiprotein complex data as hypergraphs identifies some different proteins as ‘hubs’ than its representing graph. In Subsection 3.3, we compare the identified degrees with gene-essentiality information.

**Figure 4:**
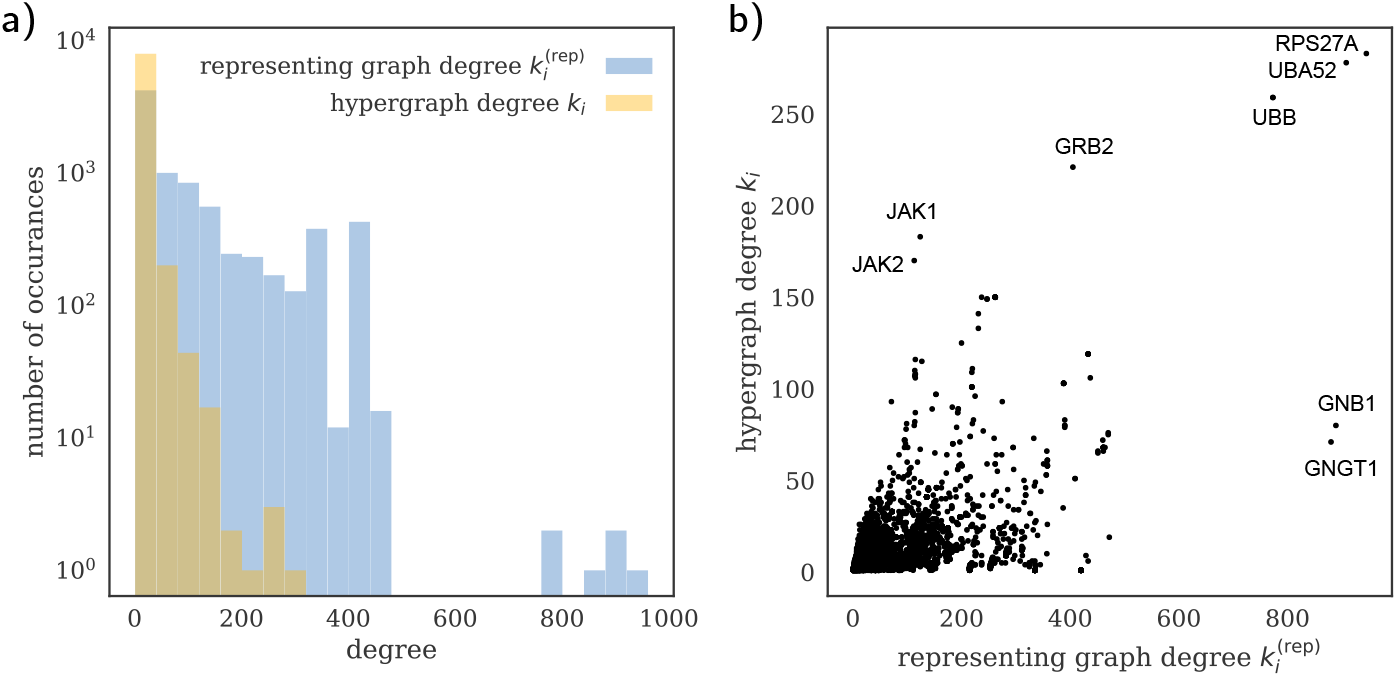
(a) Distribution of degrees in hypergraph and its representing graph. (b) Scatterplot of the hypergraph degree and the degree in the representing graph.

**Figure 5:**
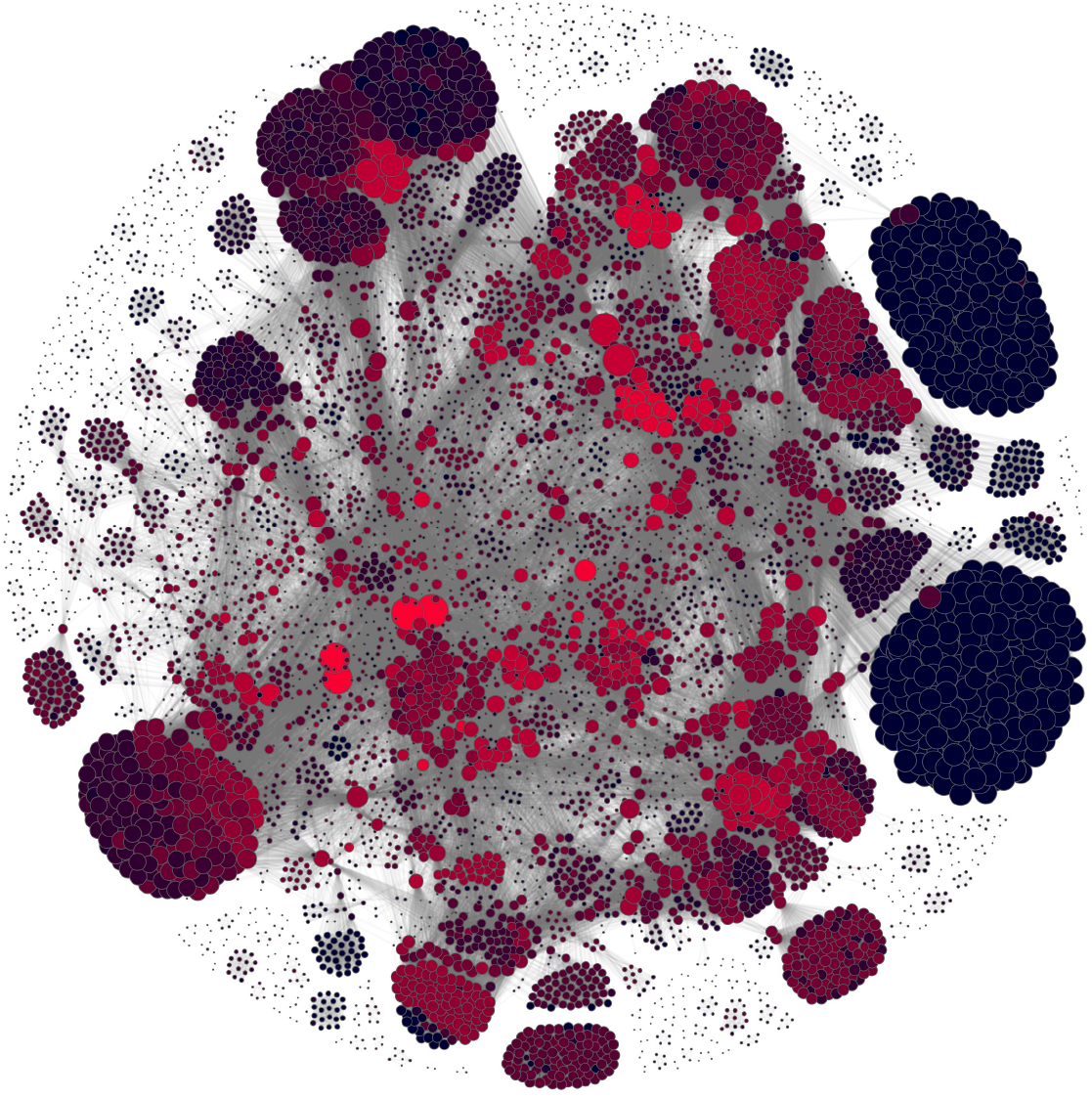
Force-directed layout of the representing graph. The size of nodes is in proportion to the representing graph degree 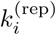 and the colour indicates the hypergraph degree 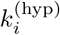 from low (blue) to high (red). Accordingly, large, blue nodes indicate proteins with a high representing graph degree and a low hypergraph degree. Illustration created with Netwulf [41].

### 3.3 Identifying essential proteins and protein complexes

One of the prominent applications of protein-protein interaction networks is the identification of essential proteins (i.e., proteins without which an organism cannot survive). The degree of proteins has been suggested as a way to predict essentiality [5]. We now assess whether the degree in the hypergraph is also able to predict the essentiality of proteins.

For this task, we compare the average degree of proteins expressed by essential genes with the average degree of proteins expressed by genes that are not essential or conditionally essential (see Table 2). We observe that in the hypergraph essential proteins have a higher mean degree than non-essential proteins and conditionally essential proteins lie between those. This indicates that the more multiprotein complexes proteins participate in, the more functionally important they are. We use a χ^2^-test to investigate the null hypothesis that essential and non-essential proteins belong to the same population. We obtain a χ^2^ ≈ 1459 with one degree of freedom (p-value < 10^−5^) and reject the null hypothesis, which is strong evidence that hypergraph degrees of proteins can predict essentaility of genes.

**Table 2:** The mean degree ⟨*k*_*i*_⟩ of nodes which connect to essential proteins (i.e., proteins expressed by essential genes), to conditionally essential proteins, and to non-essential proteins in the hypergraph and its representing graph.

| mean degree | hypergraph | representing graph |
| --- | --- | --- |
| essential | 14.06 | 85.48 |
| conditional | 9.87 | 77.45 |
| non-essential | 6.444 | 110.89 |

In the representing graph, we do not observe that essential proteins tend to have a higher degree than non-essential proteins. This occurs because essential proteins tend to be connected to hyperedges of low cardinality. Our results indicate that the hypergraph representation is more fruitful than the representing graph for this application.

A further advantage of the hypergraph in comparison with the representing graph is that we can associate a protein complex with each hyperedge. While there is no information available whether a certain protein complex is essential, we may infer whether a complex is potentially essential from protein-essentiality data by asserting that only a multiprotein complex that has at least one composing protein that is essential may also be essential. In total, 811 out of 6688 complexes have at least one essential protein associated with them and are therefore *potentially essential* protein complexes (see Fig. 6). The Spearman correlation between the number *c*_essential_ of essential proteins connected by a hyperedge and the hyperedge’s cardinality *c* is 0.35 (*p* < 10^−193^). This is in accordance with earlier findings on yeast that larger complexes tend to be more essential [42].

**Figure 6:**
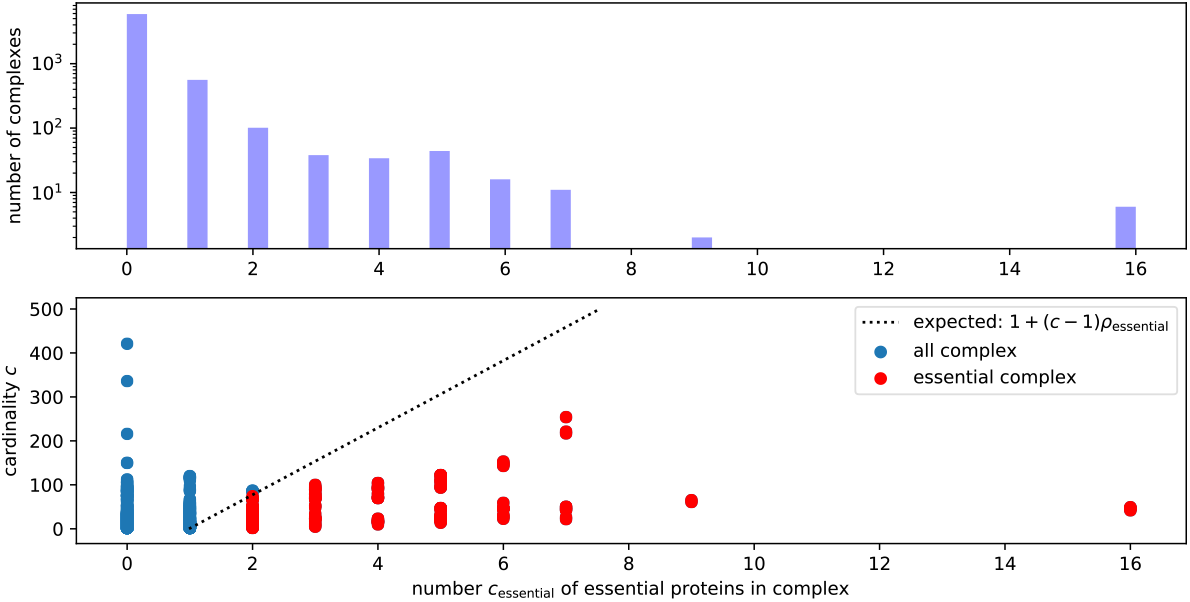
(Upper Panel) The distribution of the number *c*_essential_ of essential proteins in multiprotein complexes. (Lower Panel) The number *c*_essential_ of essential proteins versus the cardinality *c* of the complexes. We highlight complexes that have more essential proteins than expected under a random null model (dashed line) in red.

To test which of the protein complexes consist of more essential proteins than expected by chance, we construct a random null model: For each essential hyperedge of cardinality *c* we fix one essential protein in the hyperedge and sample *c* − 1 proteins. Out of a total of 8243, proteins 108 are essential, which gives a density of essential proteins of *ρ*_essential_ ≈ 0.013. Assuming that we pick *c* − 1 proteins at random with replacement, independently of each other, each pick would have probability *ρ*_essential_ ≈ 0.013 of being essential. Therefore, under this model, the number of essential proteins in an essential complex would follow a shifted binomial distribution with an expectation value of *E*(*c*) = 1 + (*c* − 1)*ρ*_essential_, which we show as dashed line in the lower panel of Fig. 6. In our data set, 246 out of 6688 protein complexes are essential and shown as red disks in Fig. 6. When comparing the fraction *c*_essential_/*c* of essential proteins to the size *c* of the complex, we observe an anticorrelation (see Figure in SI B). This indicates that larger complexes tend to be more essential because they are larger and not because they have a higher density of essential proteins.

Overall, this analysis indicates that the hypergraph degree is in better agreement with gene-essentiality information than the representing graph. Additionally, the hypergraph allows us to statistically investigate the essentiality of protein complexes.

### 3.4 Hierarchy coefficient

The results above indicate that the hypergraph contains biological signal and conveys different information than the representing graph. Next, we find that the representing graph may have a hierarchical structure which arises solely from the translation of hyperedges into simple edges.

For many complex networks, it has been reported that the local clustering coefficient follows the degree in a power law, which has been interpreted as a sign of a hierarchical organisation [26]. First, we compute the local clustering for the hypergraph and for the representing graph. Fig. 7 shows that on average, the local clustering *C*_*i*_ is much larger in the graph representation than in the hypergraph. The average local clustering ⟨*C*_*i*_⟩_graph_ ≈ 0.810 in the graph is an order of magnitude larger than of the hypergraph ⟨*C*_*i*_⟩_hypergraph_ ≈ 0.078. We find that the local clustering between both graphs is anticorrelated (Spearman correlation −0.49). This is plausible because some high-cardinality hyperedges connect many different proteins, which leads to a local hypergraph clustering close to 0 but a local graph clustering close to 1 (see right panel of Fig. 7 for an extreme example). This indicates that constructing a representing graph from a hypergraph inflates the clustering coefficient for nodes incident to such high-cardinality hyperedges.

**Figure 7:**
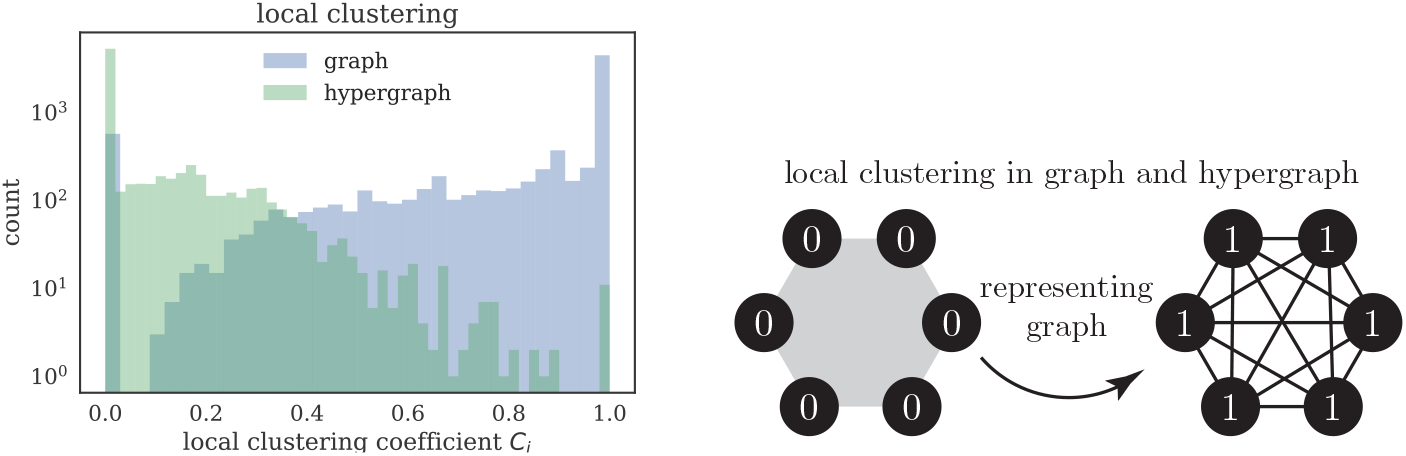
(Left Panel) The distribution of local clustering *C*_*i*_ differs strongly between hypergraph and graph. (Right Panel) An example showing that replacing hyperedges with high cardinality with a *c*-clique may inflate the local clustering from zero to one.

To explore the hierarchical organisation for hypergraphs, we investigate the relationship between clustering and degree for the hypergraph and the representing graph (see Fig. 8). For the hypergraph, we estimate the power-law exponent *β*_hyp_ ≈ 0.001 (p-value 0.899) and thus do not observe a hierarchical organisation. For the representing graph, however, we find *β*_rep_ ≈ 0.07 (p-value 4 × 10^−9^) and thus observe an organisation that appears to be ‘hierarchical’ in the sense of [26].

**Figure 8:**
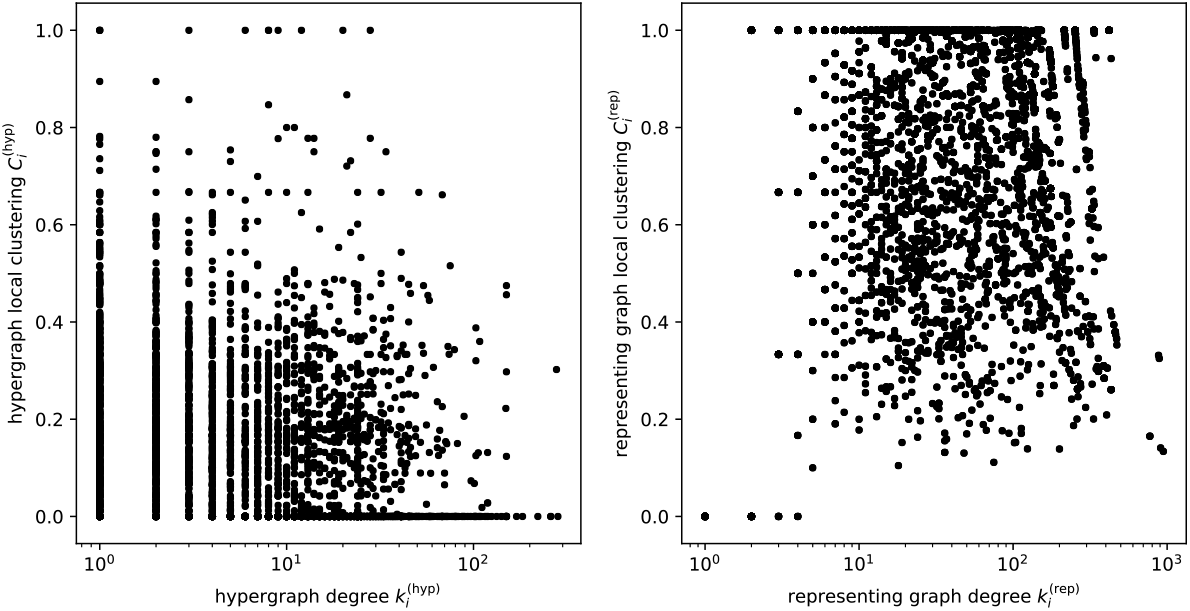
(Left Panel) Degree and local clustering for all nodes in the hyper-graph. (Right Panel) Degree and local clustering for all nodes in the representing graph.

This indicates that hypergraphs that themselves do not show a statistically significant relationship between local clustering *C*_*i*_ and degree *k*_*i*_ can show a statistically significant relationship after being translated into their representing graph. This behaviour can be explained by the pairwise projection procedure in Fig. 7b: hyperedges of cardinality *c* that do not intersect with any other edges are replaced with a *c*-clique in the representing graph. This creates *c* nodes with degree *k*_*i*_ = *c* − 1 and clustering coefficient *C*_*i*_ = 1. As we have many hyperedges with only small overlap with other hyperedges, we obtain many of such nodes in the representing graph, which creates an apparent hierarchical organisation.

## 4 Discussion

Multiprotein complexes are biological polyadic interactions between proteins that can be represented by hypergraphs. In this paper, we have used a hy-pergraph representation of the data and using two null models for hypergraphs have found that the data hypergraph contains signal beyond their cardinality sequence **c** and their degree sequence **k**. Similar results are well-established for protein interaction graphs but have not been tested for multiprotein complex hypergraphs [43]. By projecting the hypergraph into a graph representation, we illustrate that this simplification reveals different degree-rankings, which indicates that using both mathematical structures may reveal complementary information. In our test on human data, the hypergraph representation revealed a stronger correlation with gene-essentiality information than the representing graph. We then estimated the essentiality of protein complexes by comparing it with a null model and found that larger complexes tend to be more essential. In future work, one could investigate whether other hypergraph centralities (e.g., an eigenvector-based centrality [44]) are in even better agreement with essentiality.

Using an established definition of local clustering coefficient in hypergraphs, we defined the hierarchy coefficient for hypergraphs. We then showed that a pairwise graph may appear to show a hierarchical organisation while the hypergraph does not. As graphs are often constructed from polyadic interaction data, this finding reveals that such results might occur through the projection process and not the biological systems themselves.

In this study, we have demonstrated that hypergraphs are a fruitful representation of higher-order interactions between proteins. We did, however, ignore the stoichiometry (i.e., the number of proteins of a certain type that are involved in a complex). The investigation of a mathematical structure that incorporates such information might be a fruitful extension to our work. Furthermore, one could consider the different role (e.g., catalyst) that proteins have in chemical reactions and investigate them as *annotated hypergraphs* [45].

The formation of stable protein complexes investigated in this study is just one way in which proteins interact with each other. There exist many *transient* protein interactions that form and break on shorter time scales but are nevertheless of crucial biological importance [46]. An integrative analysis of pairwise protein interaction sources (e.g., BIOGRID) with multiprotein-complex data may reveal a more nuanced picture of the cellular processes than either data set on their own.

Hypergraphs and their null models might be used to analyse other data sets. Among them are association-based data (e.g., ingredient-product networks [47, 48], authorship networks [49], company-directorate networks [50]) or social networks in which higher-order interactions have been shown to be prominent [51, 52]. The tools in this manuscript could be used to investigate such data as hypergraphs and so reveal organisational principles beyond their pairwise interactions.

## 5 Data and code availability

We make the code and data available on GitHub under https://github.com/floklimm/hypergraph.

## 6 Funding

F.K. was supported by the Engineering and Physical Sciences Research Council (funding references EP/R513295/1 and EP/N014529/1). G.R. was supported in part by the EPSRC grant New Approaches to Data Science: Application Driven Topological Data Analysis EP/R018472/1.

## 7 Acknowledgements

We thank Christopher Koch, Christina Goldschmidt, and Phil Chodrow for fruitful discussions.

## Supplementary Information

### A Dual graph

#### A.1 Definition

The *dual graph* 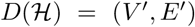 (also called *line graph*) of a hypergraph 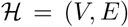 is the graph whose vertex set *V*′ is the set of hyperedges of the hypergraph with edges between them if the hyperedges have at least one node in common (i.e., *V*′ = *E* and (*e*_*i*_, *e*_*j*_) ∈ *E*′ ↔ *e*_*i*_ ∩ *e*_*j*_ ≠ ∅). In Fig. 9, we show an example of the dual graph constructed from a hypergraph.

**Figure 9:**
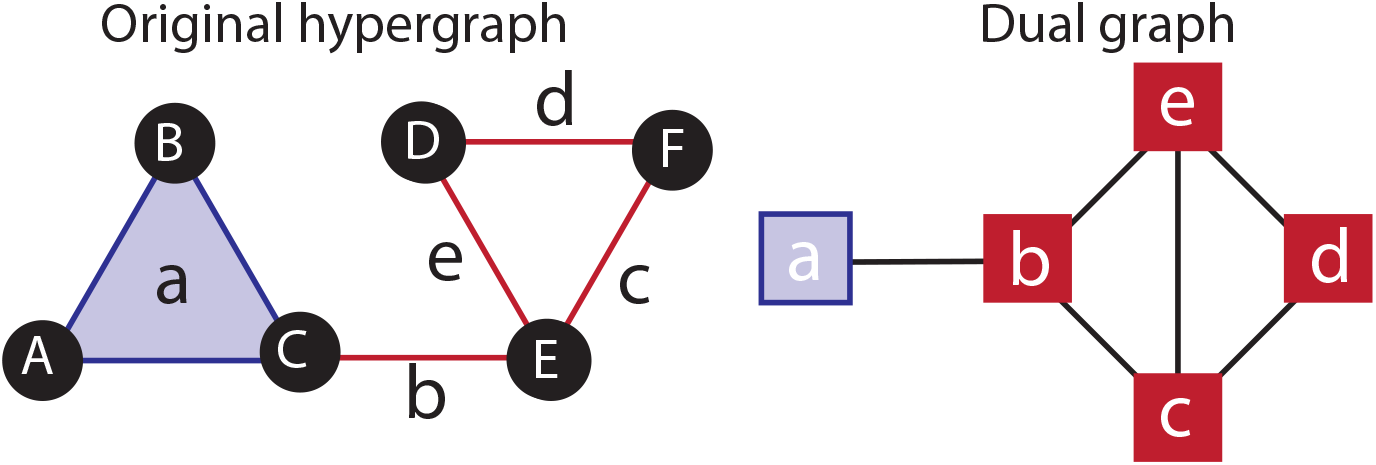
Example of a *dual graph* constructed from a hypergraph. Each hyperedge is represented by a node. These nodes are connected if the hyperedges share a node.

#### A.2 Results for the dual graph

In Fig. 10 we compare the hypergraph with its dual graph, keeping in mind that in the dual graph 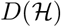 the nodes represent the edges of the hypergraph 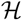. Therefore, we compare the degree 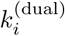 of dual graph nodes *V*′ with the cardinality of the hyperedges *E* in the original hypergraph (see Fig. 10). There are 1717 hyperedges with minimum cardinality *c*_min_ = 2. This is the cardinality that occurs most often. The mean cardinality is ⟨*c_e_*⟩ ≈ 10.12. and the mean degree is 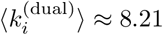.

**Figure 10:**
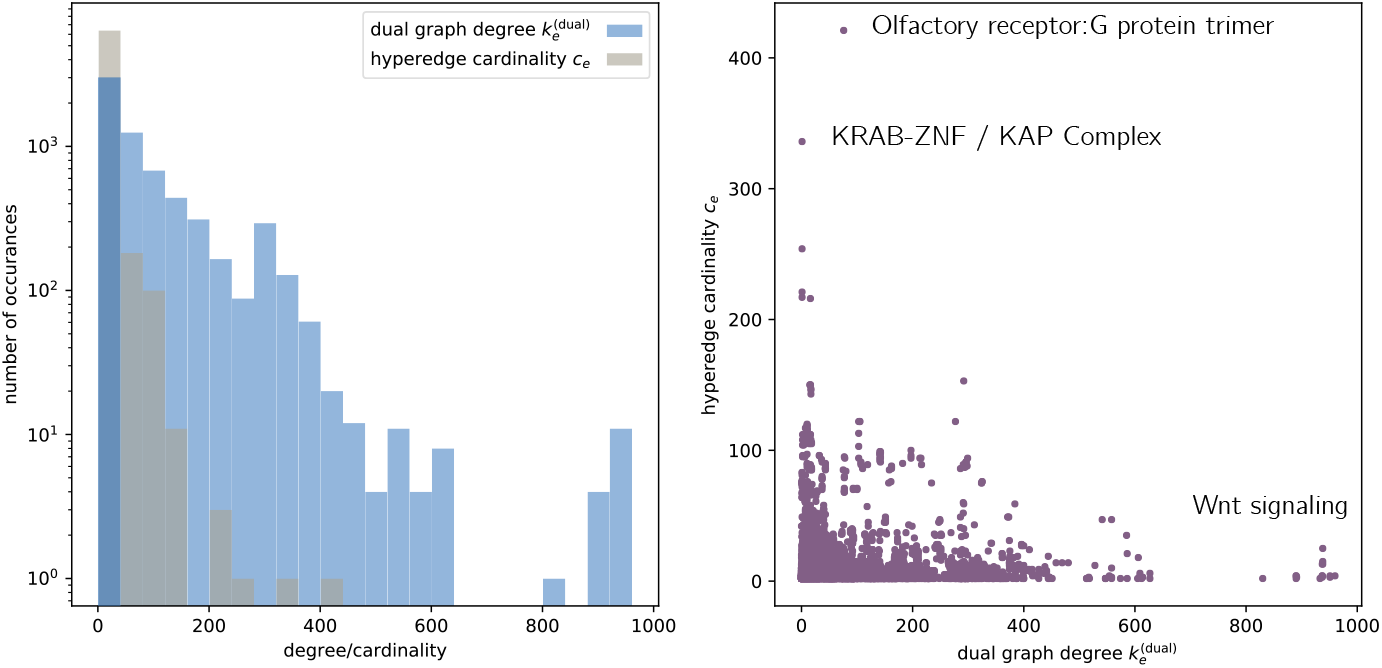
(Left) Distribution of *c*_*e*_ cardinality of hyperedges in *H* and the degrees of the associated nodes in the dual graph *D*(*H*) (Right) Scatterplot of these two.

In the dual graph 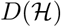, we also investigate the degrees of nodes and compare it with the cardinality of the hyperedges *E* in the original hypergraph 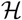. The Pearson correlation is −0.03 and the Spearman correlation of −0.03, indicating that the size of the complex and the degree of its associated node in the dual graph are almost uncorrelated.

The node in 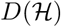 with the highest degree 960 represents a Wnt complex and has a cardinality of 119. We observe that there are multiple other nodes that have a slightly lower degree and similar cardinality. All of these nodes in 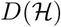 represent complexes that are also involved in the Wnt signaling pathway and are active in the clathrin-coated endocytic vesicle membrane, which plays a critical role in the Wnt signaling pathway [53]. This pathway itself is crucial for stem cell development and disease progression [54]. The complex with the highest cardinality of 421 is the ‘Olfactory receptor-G protein trimer’. It has a degree of 83. The second highest cardinality has the ‘KRAB-ZNF / KAP Complex’ with a cardinality of 336 and a degree of 6.

This investigation illustrates that degree of the dual graph and cardinality of the protein complexes identify distinct protein as high ranked. Both approaches reveal protein complexes of crucial cellular function and are therefore fruitful strategies to investigate cellular hypergraphs.

### B Fraction of essential proteins in multiprotein complexes

**Figure 11:**
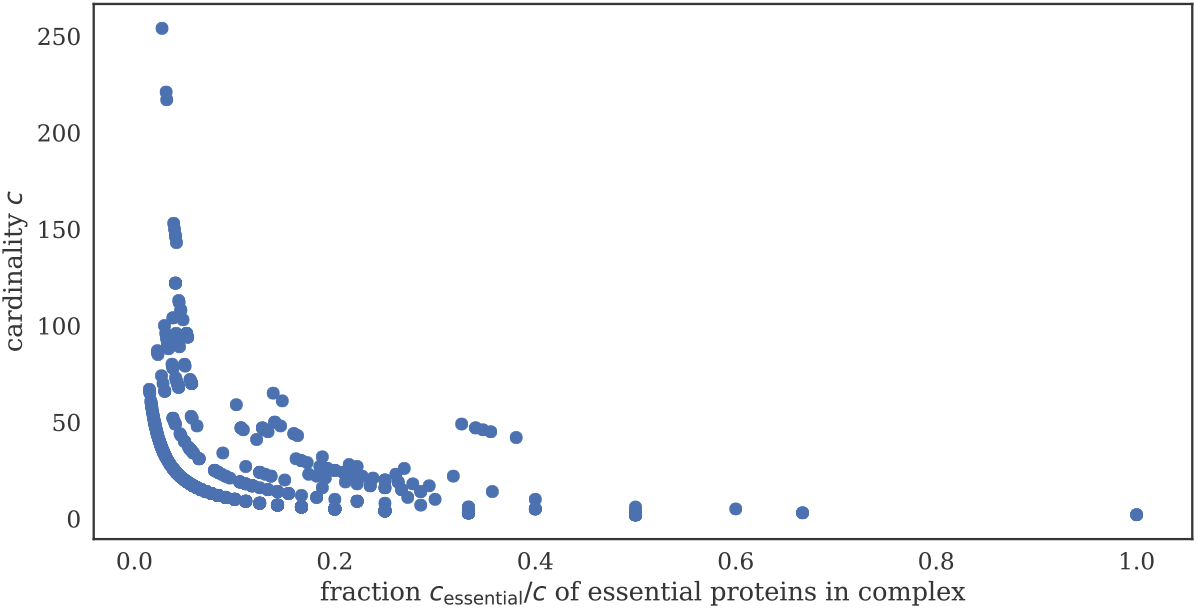
The fraction *c*_essential_/*c* of essential proteins in complexes, in dependence of the cardinality *c*. The two are anticorrelated, i.e., larger protein complexes tend to have a smaller fraction of essential proteins.

### C Distribution of hypergraph measures for the null models

**Figure 12:**
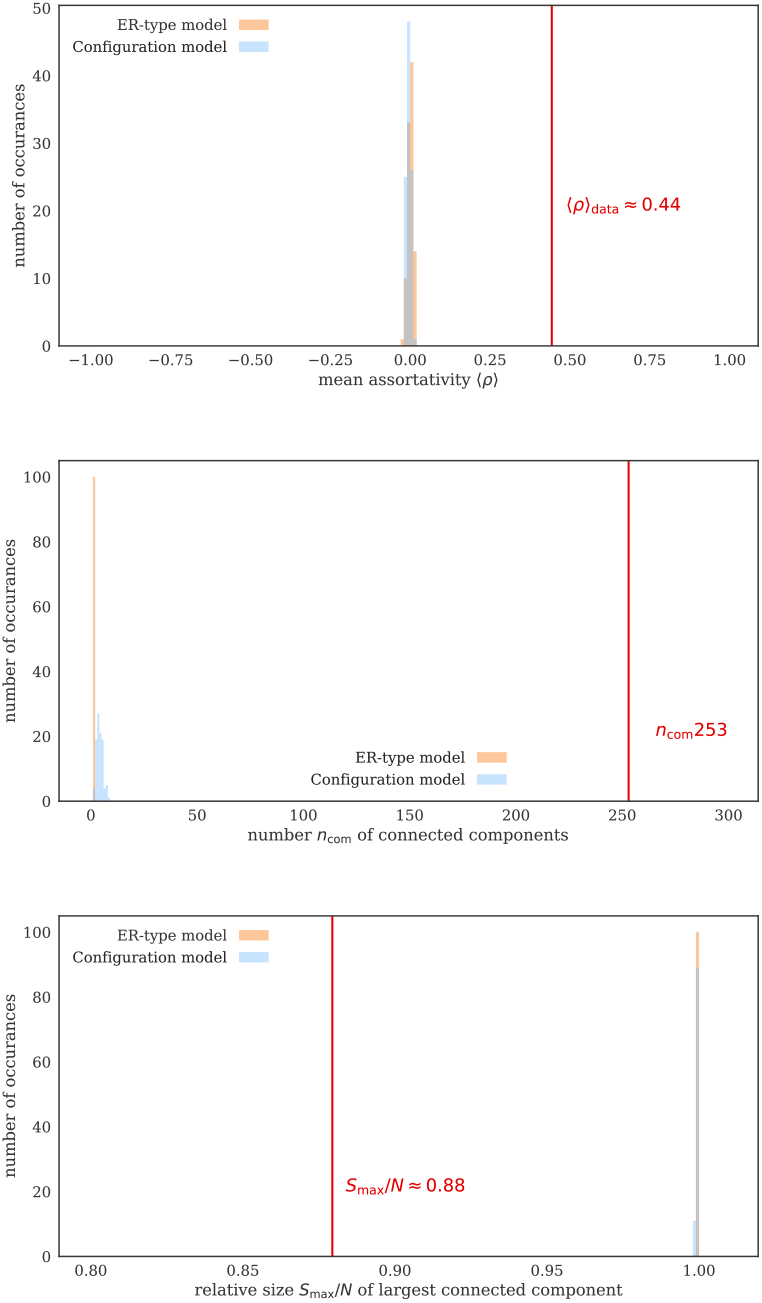
The distribution of mean assortativity ⟨*ρ*⟩, number *n*_com_ of components, and relative size *S*_max_/*N* of the largest component for the ER-type model (orange) and the configuration model (blue) for 100 realisations. The mean assortativity ⟨*ρ*⟩_data_ ≈ 0.44 of the protein hypergraph (red vertical line) is clearly larger than for these null models. The number *n*_com_ = 253 of components is also larger for the protein hypergraph. The relative size *S*_max_/*N* ≈ 0.88 of the largest component is smaller.

**Figure 13:**
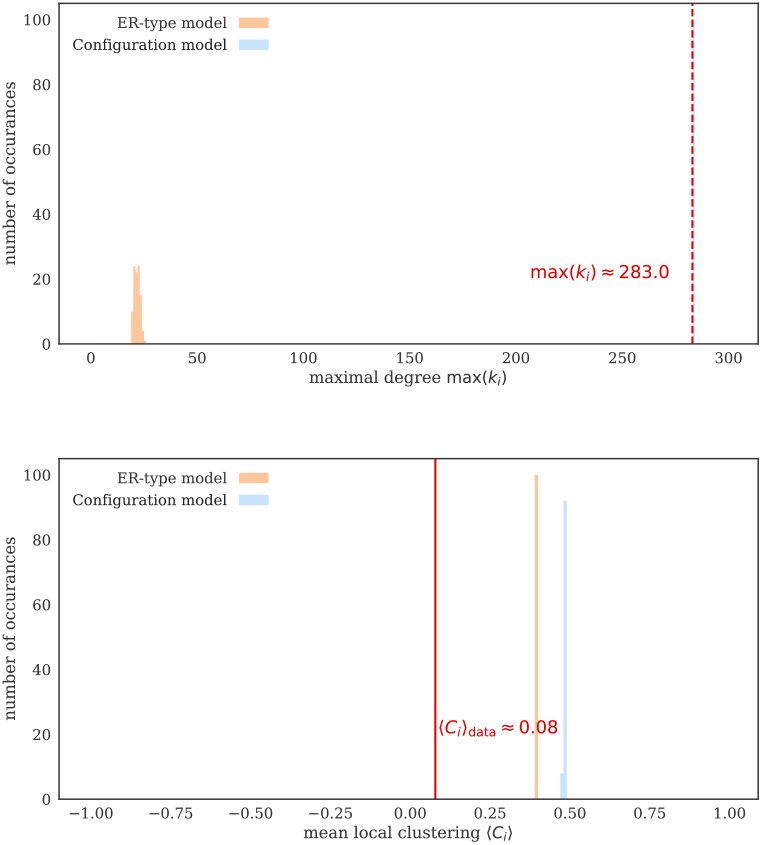
The distribution of the maximal degree max (*k*_*i*_) and mean local clustering ⟨*C*_*i*_⟩ for the ER-type model (orange) and the configuration model (blue) for 100 realisations. The maximal degree is (by construction) the same for the configuration model as for the protein hypergraph. The ER-type model has a much smaller maximum degree. The mean local clustering ⟨*C*_*i*_⟩ ≈ 0.8 is smaller for the data than for the null models.

## Footnotes

1 The convex hull of any subset of the *n* + 1 points that define a *n*-simplex is called a *face*. A simplicial complex 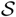 is a set of simplices in which every face of a simplex is also in 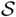 [19]. This property is called *set inclusion*.

## Notes

https://github.com/floklimm/hypergraph

